# Cultured Mesenchymal Cells from Nasal Turbinate as a Cellular Model of the Neurodevelopmental Component of Schizophrenia Etiology

**DOI:** 10.1101/2023.03.28.534295

**Authors:** Victoria Sook Keng Tung, Fasil Mathews, Marina Boruk, Gabrielle Suppa, Robert Foronjy, Michele Pato, Carlos Pato, James A. Knowles, Oleg V. Evgrafov

**Author notes:** **Correspondence:** Oleg V. Evgrafov.

## Abstract

Study of the neurodevelopmental molecular mechanisms of schizophrenia requires the development of adequate biological models such as patient-derived cells and their derivatives. We previously used cell lines with neural progenitor properties (CNON) derived from superior or middle turbinates of patients with schizophrenia and control groups to study gene expression specific to schizophrenia.

In this study, we compared single cell-RNA seq data from two CNON cell lines, one derived from an individual with schizophrenia (SCZ) and the other from a control group, with two biopsy samples from the middle turbinate (MT), also from an individual with SCZ and a control. In addition, we compared our data with previously published data from olfactory neuroepithelium (1). Our data demonstrated that CNON originated from a single cell type which is present both in middle turbinate and olfactory neuroepithelium. CNON express multiple markers of mesenchymal cells. In order to define relatedness of CNON to the developing human brain, we also compared CNON datasets with scRNA-seq data of embryonic brain (2) and found that the expression profile of CNON very closely matched one of the cell types in the embryonic brain. Finally, we evaluated differences between SCZ and control samples to assess usability and potential benefits of using single cell RNA-seq of CNON to study etiology of schizophrenia.

## Introduction

Schizophrenia (SCZ) is a brain disease with a complex etiology that commonly develops during adolescence or early adulthood. It is widely considered that alterations in brain development play a significant role in the etiology of the disease (3). While post-mortem brain samples can be used to investigate epigenetic, transcriptomic, and proteomic alterations in SCZ patients’ brains, disease-specific alterations in neuronal cells during embryonic and fetal brain development requires the use of cellular models.

Patient-derived induced pluripotent stem cells (iPSCs) and iPSC-derived neural progenitors or neurons are often used as cellular models to study the neurodevelopmental aspects of SCZ and other disorders. However, iPSCs-derived cellular models represent *in vitro* induced neurodevelopment from cells regaining stemness properties after genetic alterations. They more likely reproduce very early events of neurodevelopment and may not capture specifics of neurodevelopment in complex biological system in sufficient detail, especially at later stages.

An alternative approach is to use cells derived from olfactory neuroepithelium (ON) where neurogenesis occurs throughout life. The brain and ON develop from neighboring ectoderm regions with some contributions from the neural crest (4,5). ON develops from ectoderm during the 5th week of post-fertilization age (PFA) (4). Although the brain undergoes a more advanced developmental process, we may expect that the brain at *early* stages of development will still share some of the cell types of ON due to a common origin from neuroepithelium. While there is no specific data about the degree of relatedness between developed nasal turbinate and embryonic brain, the ability of ON to produce neuronal cells suggests that the similarity between them could be substantial, at least in some cell types.

The hypothesis that ON could serve as a model to study SCZ is further supported by high correlation between SCZ and anosmia, affecting neuronal functions in brain and ON respectively and findings of dysregulation of olfactory neuron lineages in SCZ (6).

Several cellular models have been developed using different protocols (7–12). In our previous studies, we used a protocol for developing cell cultures from ON which was originally proposed by Wolozin et al. (7). The key element of this protocol is to cover small pieces of ON with Matrigel and propagate only those cells that penetrate through the gel (see protocol details in (13)). We named these cells CNON, Cultured Neuronal Cells derived from Olfactory Neuroepithelium, and considered them neural progenitors, although their ability to differentiate into neurons *in vivo* has not been proven. It was subsequently discovered that the same cell type could be generated from both superior and middle turbinate (14).

We have previously developed CNON from 256 individuals including 144 patients with schizophrenia (SCZ) and demonstrate the robustness of CNON development and consistency of expression profiles during growth in culture and between individuals (13). Using single-cell transcriptomics, we identified a cell type in middle turbinate (MT) with an expression profile corresponding to CNON (15), confirming that CNON is not a mixture of cell types but instead a single cell type with a specific gene expression profile. These properties resulted in lower biological noise as compared to cellular models with higher heterogeneity. It warrants higher statistical power and requires a smaller number of samples to identify the signal. With advances in single-cell transcriptomics, it becomes possible to assess the similarity of CNON to cells in the brain in more detail and evaluate the relevance of this model to study brain disorders.

## Materials and Methods

### Biopsy collection and sample preparation

Biopsies were obtained from patients without any history of sinonasal disease or surgery or immunocompromise. Tissue samples were obtained from the superior-medial region of the head of the middle turbinate; laterality was determined by ease of access. The mucosa was anesthetized and decongested with topical 1% lidocaine and 0.05% oxymetazoline. After 5 minutes, 0.3 ml of 1% lidocaine with 1:100,000 epinephrine was injected into the targeted mucosal site under visualization. 2mm cupped forceps were used to obtain biopsy specimens. These samples were immediately transported to the lab in Leibovitz’s L-15 Medium (ThermoFisher) supplemented with Antibiotic-Antimycotic solution (ThermoFisher) and processed to prepare single-cell suspension. Biopsy pieces were minced with two scalpels in a Petri dish with a small amount of cold Hank’s buffer without magnesium or calcium to prevent drying and then transferred to a 1.5 ml Eppendorf tube and washed twice with 1 ml cold Hank’s buffer. Cells were dissociated in 250 μl of 0.25% Trypsin-0.02% EDTA (VWR) at 37°C while shaking at 500 rpm. After 10 minutes, the suspension was mixed by pipetting and returned to thermomixer for another 10 minutes. After another mixing, the cells were sedimented by centrifugation at 300 relative centrifugal force for 5 minutes, resuspended in 500 μl of cold Hank’s solution (ThermoFisher) with BSA, filtered using 40 μm FLOWMI cell strainer (Bel-Art) cell suspension into a 1.5 ml Eppendorf tube, centrifuged again, and finally resuspended in 50 μl of Hank’s with 0.04%BSA (ThermoFisher).

### CNON cell culture

Protocols for developing CNON cell cultures from middle or superior turbinate biopsies have been previously described (13). In brief, each biopsy sample was dissected into 3-4 pieces approximately 1 mm^3^ in size, and each piece was placed onto the surface of a 60 mm tissue culture dish coated with 25 μl of Matrigel Basement Membrane (Corning) reconstituted in F12 Coon’s medium (Sigma), and then covered by 15 μl of full-strength Matrigel. After Matrigel gelatinizes, 5 ml of medium 4506 (16) was added. Medium 4506 is based on F12 Coon’s medium (Sigma) supplemented with 6% of FBS (KSE scientific), 5 ug/ml human Gibco transferrin (Fisher Scientific), 1 ug/ml human insulin (Sigma), 10 nM hydrocortisone (Sigma), 2.5 ng/ml sodium selenite (Sigma), 40 pg/ml thyroxine (Sigma), 1% Gibco Antibiotic-Antimycotic (Fisher Scientific), 150 μg /ml Bovine hypothalamus extract (Millipore-Sigma) and 50 μg/ml Bovine pituitary extract (Sigma). Within 1-4 weeks of incubation, neuronal cells CNON cells were observed to grow out of the embedded pieces of tissue. Due to unique ability to grow through Matrigel, neural progenitors often populate large areas without presence of other cell types. Outgrown cells with a neuronal phenotype were then physically isolated using cloning cylinders and dislodged using 0.25% Trypsin-0.02% EDTA (VWR) and transferred into a new Petri dish for further cultivation.

### Single cell preparation from CNON

Cells at ~80% monolayer on a 6 cm Petri dish were dislodged with 1 ml of 0.25% Trypsin-0.02%EDTA solution, and 3 ml 4506 culture medium was added to stop digestion. Cells were then gently and thoroughly mixed to break up clumps of cells, spun at 300 rcf for 5 min, and resuspended in a 3 ml culture medium. After gentle mixing by pipetting, the cell suspension was filtered using a 40 μm Flowmi™ cell strainer into a 15 ml tube. Cells were washed with 1x DPBS (ThermoFisher) with 0.04% BSA, then 1X PBS (ThermoFisher) with 0.04% BSA, re-suspended in 500 ul of PBS, and filtered using Flowmi™ Tip Strainer. After counting, cell concentration was adjusted to 700 cells/μl.

### scRNA-seq

The concentration and viability of cells were determined using a hemocytometer and Trypan blue. After counting, single-cell libraries were prepared according to the 10x Genomics protocol CG000183 on Chromium controller (10x Genomics) and sequenced on NovaSeq6000 as paired end 28 + 90 bp reads plus two indexing reads.

### scRNA-seq analysis

Raw sequencing data were processed using bcl2fastq2 v2.20 to convert BCL files to fastq files while simultaneously demultiplexing. Fastq files were processed using *TrimGalore* v. 0.6.5 to automate quality, adapter trimming, and perform quality control.

We used Cell Ranger v. 6.1.2 (10x Genomics) to generate raw gene-barcode matrices from the reads, which were aligned to the **GRCh38 Ensembl v93**-annotated genome. The following analysis was done using R package Seurat v.4.10 (17). First, cells with low number of detected RNA molecules or genes were removed from the data and gene expressions were normalized and scaled using *Sctranform*. We then applied a graph-based clustering algorithm in order to group cells into distinct clusters.

To account for any potential variability due to cell-cycle phase, we calculated cell-cycle phase scores using Seurat’s built-in lists of cell-cycle genes. We then regressed out these scores from the dataset during normalization in order to reduce the effect of cell cycle heterogeneity on our analysis using Seurat algorithm (**https://satijalab.org/seurat/articles/cell_cycle_vignette.html**). After analysis of cluster trees (*clustree*) we selected an optimal resolution parameter (0.5 for MT studies) for cluster analysis. We annotated the clusters using known markers and data from relevant single-cell studies, and generated UMAP plots for visualization.

To identify genes that are differentially expressed between the cell clusters in scRNA-seq data, we used the *FindAllMarkers(*) function in Seurat. This function compares the expression of each gene in each cluster using a Wilcoxon rank-sum test and returns a list of differentially expressed genes. We set the min.pct argument to 0.25 (to include only genes with at least 25% difference in mean expression) and the *logfc.threshold* argument to 0.25 (log fold change of at least 0.25 between the clusters).

To visualize the DE results, we used the *DotPlot*() function in Seurat, which plots the fold change in gene expression using the color scale and the size of the circle as percentage of cells expressing the gene. We also created a table of these DE genes using the data from the dot plot. This table allowed us to more easily review and analyze the DE genes identified by the *FindAllMarkers(*) function.

To compare expression profiles of CNON cells with single cell data from tissues (middle turbinate, olfactory neuroepithelium and embryonic brain), we performed reference mapping using Seurat, annotating the query CNON datasets on a cell-type labeled reference datasets and evaluating the mapping of predicted cell-type annotations in the query datasets to the reference dataset using the *TransferData(*) function.

ScRNA-seq data from embryonic brain was obtained from the UCSC Cell Browser (matrixes from combined samples) and the NeMO repository (https://assets.nemoarchive.org/dat-0rsydy7) where individual sample matrixes were available.

ScRNA-seq data from olfactory neuroepithelium was obtained from Gene Expression Omnibus under accession code **GSE139522**.

## Results

To assess potential utility of cultured cells derived from nasal turbinates to study the developmental mechanisms of schizophrenia, we performed scRNA-seq of cultures derived from a patient with SCZ (CNON-SCZ) and from an individual from the control group (CNON-CTRL). To identify parental cell types, we compared this with scRNA-seq data from two middle turbinate biopsy samples, taken from a patient with SCZ (MT-SCZ) and control (MT-CTRL). Lastly, CNON datasets were compared with single cell transcriptome data from the embryonic brain (2).

For comparison we employed *reference mapping* from Seurat, which mapped query datasets (CNON) to reference datasets, which were either MT or embryonic brain data. Transcriptome of each CNON cell was placed in 3-dimensional space with PC coordinates of reference dataset, and assessed if it could be assigned to any cluster of reference dataset, associated with particular cell type. The quality of assigned cell type of each query cell was evaluated by calculated *prediction score*.

Such reference mapping showed that the vast majority of CNON-CTRL cells (11741, 96%) mapped to the cluster corresponding to the Mesenchymal cell (MC) cluster of the MT-CTRL, with an average prediction score of 0.96. However, cells from MC cluster are not proliferating (do not express cell cycle-specific genes, such as *MKI67*), and the effect of cell culture genes may reduce the quality of mapping to non-proliferating cells of the same type. To mitigate the effects of CNON proliferation we assigned cells with a cell-cycle score based on the expression of canonical cell-cycle genes, applied these scores to model the relationship between gene expression and cell-cycle score and removed this cell-cycle phase variability from our data using regression. These procedures can be performed using functions in Seurat. Indeed, regressing cell-cycle score resulted in accurate mapping of all CNON-CTRL cells to MC cluster with a perfect prediction score of 1 (**Table 1**).

**Table 1.**
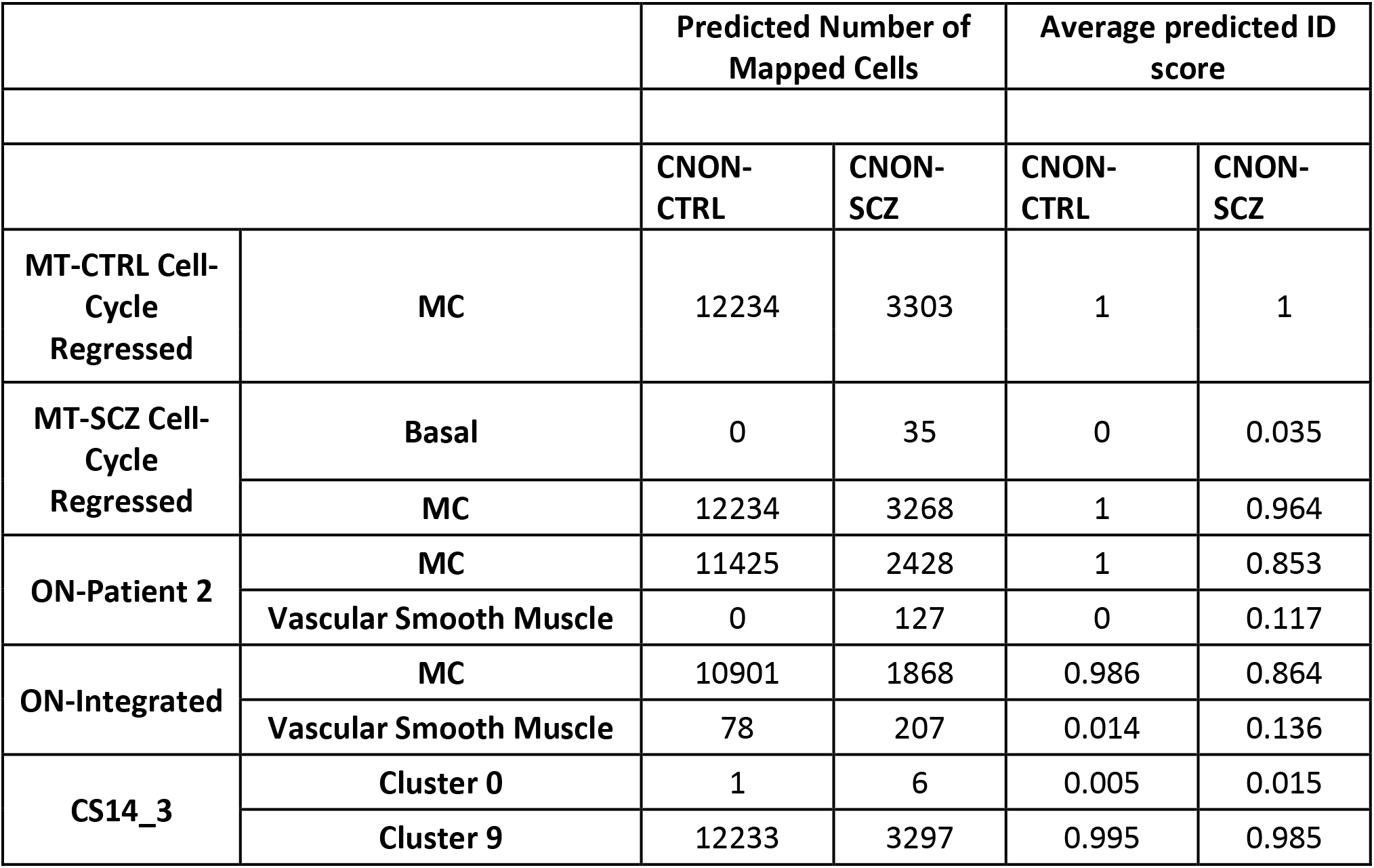
Mapping CNON cells onto various reference datasets. Number of cells from CNON-CTRL and CNON-SCZ mapped onto different cell types or clusters found in MT-CTRL (after cell-cycle score regression), MT-SCZ (after cell-cycle score regression), CS14_3, ON-Patient 2, and ON-Integrated, and average predicted identity scores for each cell cluster mapping.

**Figure 1a and b** show results of mapping of CNON data from both samples onto two MT datasets after Uniform Manifold Approximation and Projection (UMAP) for dimension reduction (18); all six figures were presented using the same axes and on the same scale. Cell types in MT-CTRL were annotated based on marker genes specific for every cluster. Such genes were identified for each cluster by differentially gene expression analysis with all other clusters. The heatmap (**Figure 1g**) of MT-CTRL reveals a clear distinction between various cell types. While **Figure 1**a and **Figure 1b** demonstrate allocation of CNON to cluster MC in UMAP coordinates, **Table 1** provides numbers of cells mapped to specific cluster with corresponding prediction scores, supporting our previous findings that CNON developed from one cell type.

**Figure 1.**
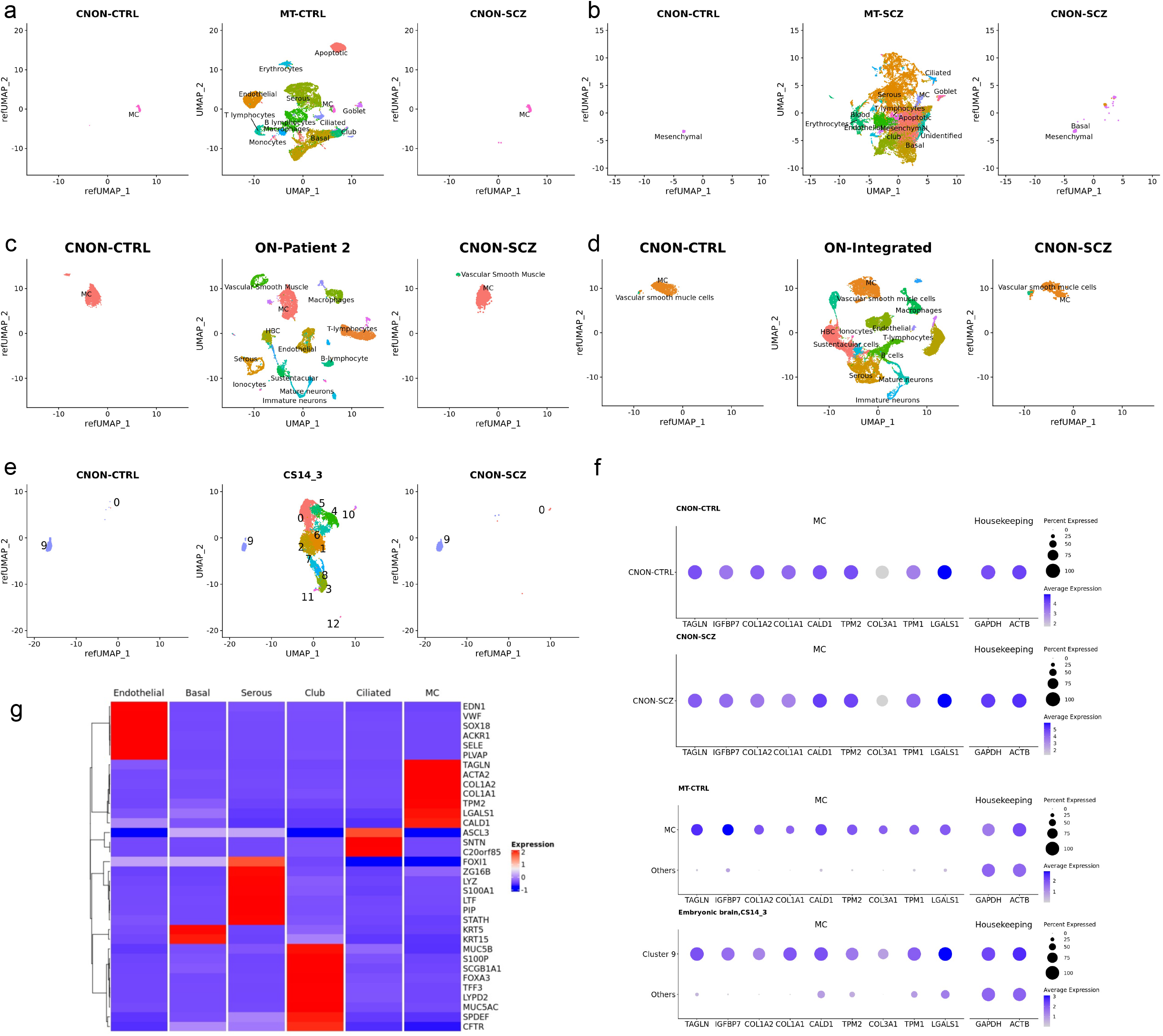
Single cell reference mapping and gene expression comparison of CNON datasets ((CNON-CTRL and CNON-SCZ) to human middle turbinate (MT-CTRL & MT-SCZ) and embryonic brain at Carnegie stage 14 (CS14_3) datasets. (A) Central panel shows UMAP dimensionality reduction plot of 21,565 human middle turbinate cells (MT-CTRL) with 13 cell types annotated. Left panel: UMAP dimensionality reduction plot of 12,234 CNON cells (CNON-CTRL). Right panel: UMAP dimensionality reduction plot of 3303 CNON cells (CNON-SCZ). All cells from both CNON-CTRL and CNON-SCZ map to the MC cluster location in MT-CTRL (B) Central panel shows UMAP dimensionality reduction plot of 28,140 human middle turbinate cells (MT-CTRL) with 13 cell types annotated. Left panel: UMAP dimensionality reduction plot of 12,234 CNON cells (CNON-CTRL), all of them mapped to MC cluster. Right panel: UMAP dimensionality reduction plot of 3303 CNON cells (CNON-SCZ). 3,268 of them map to the MC cluster location in MT-CTRL, while 35 cells mapped to Basal cluster. (C) Single cell reference mapping of CNON datasets to olfactory neuroepithelium, Patient 2. Central panel shows UMAP dimensionality reduction plot of cells from olfactory neuroepithelium of Patient 2 (1) with 13 cell types annotated. Left panel: UMAP dimensionality reduction plot of 11,425 CNON cells (CNON-CTRL); all cells mapped to Mesenchymal cell type. Right panel: UMAP dimensionality reduction plot of 2547 CNON cells (CNON-SCZ). The majority of cells (2420 CNON-SCZ cells) map to Mesenchymal cell type and 127 CNON-SCZ cells to Vascular Smooth Muscle cell clusters. (D) Single cell reference mapping of CNON datasets to olfactory neuroepithelium (integration of 4 patient samples data). Central panel shows UMAP dimensionality reduction plot of cells from olfactory neuroepithelium, integrated data from 4 patients (1) with 13 cell types annotated. Left panel: UMAP dimensionality reduction plot of 10,979 CNON cells (CNON-CTRL); 10901 cells mapped to Mesenchymal cell type, while 78 cells mapped to Vascular Smooth Muscle cells cluster. Right panel: UMAP dimensionality reduction plot of 2075 CNON cells (CNON-SCZ). The majority of cells (1,868 CNON-SCZ cells) map to Mesenchymal cell type and 207 CNON-SCZ cells to Vascular Smooth Muscle cells clusters. (E) Single Cell Reference Mapping of CNON datasets to embryonic brain (CS14_3). Central panel: UMAP dimensionality reduction plot of CS14_3 with 13 clusters. Left panel: UMAP dimensionality reduction plot of 12,234 CNON cells (CNON-CTRL); 12,233 cells mapped to to Cluster 9, 1 cell maps to cluster 0. Right panel: UMAP dimensionality reduction plot of 3303 CNON cells (CNON-SCZ). 3297cells map to cluster 9 and 6 cells map to cluster 0. (F) Average gene expression of selected cell markers in CNON. A gene considered MC marker gene if it is expressed in the MC cluster higher than in all other clusters with (a) statistical significance, (b) logFoldChange >2, and (c) expressed in at least 50% of cells of MC. MC markers: *TAGLN, COL1A2, COL1A1, CALD1, TPM2, COL3A1, TPM1*, and *LGALS1;* Housekeeping markers: *GAPDH* and *ACTB;* Neural stem cell marker: *ITGB1* are shown in CNON-CTRL, CNON-SCZ, MT-CTRL, and Embryonic brain (sample CS14_3). The size of the dot represents the percentage of cells expressing the gene, and the colors represent the average expression level of each gene.

Transcripts of MC markers are found in the vast majority of CNON cells. Meanwhile, the expression of prominent markers of other middle turbinate cell types: basal (*SERPINB3, KRT5*), endothelial (*CCL14, VWF*), serous (*DMBT1*), club (*LYPD2, SCGB1A1*), ciliated (*SNTN*), goblet cells (*MUC5B*)and ionocytes (*CFTR*) were either not present or insignificant as they were identified in less than 1% of CNON. Expression of all these genes was also low in bulk RNA-seq from CNON (**Table 2**).

**Table 2.** Expression of marker genes of major respiratory epithelial cell types in CNON (CNON-CTRL and CNON-SCZ), percentage of CNON cells and bulk CNON (average Transcripts per Million transcripts from 255 CNON samples) expressing these markers.

|  |  | CNON-CTRL |  | CNON-SCZ |  | Bulk CNON |
| --- | --- | --- | --- | --- | --- | --- |
|  |  | Average Expression | Percentage of Cells Expressing Gene | Average Expression | Percentage of Cells Expressing Gene | TPM |
| Housekeeping | ACTB | 2.456 | 100 | 2.456 | 100 | 1540.14 |
|  | GAPDH | 1.697 | 100 | 3.4082 | 100 | 1153.02 |
| Basal | SERPINB3 | 0.00375 | 0.0172 | 0 | 0 | 0.01 |
|  | KRT5 | 0 | 0 | 0.0005 | 0.303 | 0.21 |
| Endothelial | CCL14 | 0 | 0 | 0 | 0 | 0.20 |
|  | VWF | 0.0009 | 0.058 | 0.0031 | 0.182 | 0.09 |
| **Serous** | **DMBT1** | 0.0001 | 0.0007 | 0.006 | 0.394 | 0.04 |
| **Club** | **LYPD2** | 0 | 0 | 0 | 0 | 0.02 |
|  | **SCGB1A1** | 0 | 0 | 0 | 0.003 | 0.00 |
| **Ciliated** | **SNTN** | 0.0005 | 0.0327 | 0.004 | 0.272 | 0.23 |
| **Goblet** | **MUC5B** | 0 | 0 | 0.001 | 0.0606 | 0.06 |
| **Ionocytes** | **CFTR** | 0 | 0 | 0.0005 | 0.0303 | 0.07 |

We also mapped CNON data to single cell datasets from olfactory neuroepithelium (1). It contains data from four patients. We focus our investigation on reference mapping CNON cells to Patient 2, as it has the most neuronal cells per sample and we believe that it is most accurately representing olfactory neuroepithelium. The mapping result showed that CNON-CTRL exclusively mapped to a single cluster in the data of Patient 2 (**Figure 1c**) and this cluster expresses markers similar to MC cluster in MT-CTRL and MT-SCZ. The majority of cells in CNON-SCZ also map to this same cluster. Additionally, we compared our CNON cells with integrated data from all 4 patients, and most CNON cells correspond to a single cell cluster that exhibits the same gene expression profile (**Figure 1c&d**).

We then compared CNON expression profile with a large dataset of brains at several embryonic stages of development (Carnegie stages 13-22) (2). Dataset from early stages of development was chosen for our comparison as we believe that the embryonic brain was more likely to contain CNON-like cells than fetal brain at later stages of development, due to a closer relationship to a common or similar ancestors.

Comparing CNON single-cell data with scRNA-seq in this study with embryonic brain at different stages of development shows similarity with cluster 47, described by the authors in supplementary data (2). Most cells from cluster 47 were sourced from one sample designated as CS14_3, and we performed reference mapping of CNON data to this sample alone. 99.99% of CNON-CTRL (all except one cell) mapped to a single cluster 9 of CS14_3 (**Figure 1e**, **Table 1**), with an average prediction score of 0.995. This cluster corresponds to cluster 47 in the analysis of data from all embryonic samples (19). Similarly, reference mapping of a smaller dataset of CNON-SCZ resulted in 3297 cells (98.5%) annotated as cluster 9 with an average prediction score of 0.985, while 6 cells were assigned to cluster 0 with a prediction score of only 0.015 (**Table 1**).

Our findings indicate that CNON exhibits a highly comparable expression profile with one cell type found in the developing brain. In the article describing the embryonic brain dataset (2), cluster 47 was classified as “others” and it was distinctly different from explicitly classified cell types such as neurons, radial glia, neuroepithelial, intermediate progenitors, and mesenchymal cells. In sample CS14_3 this cell type accounted for about 3.5% of the total population of cells, while in other samples, including those from earlier CS13, another sample from CS14, and samples from later stages CS15, CS20 and CS22 their proportion was lower. According to our calculation these cells make up 0.87% of all cells, while the authors estimate their fraction even higher at 1.1%.

To better characterize this MC cell type, we performed gene ontology enrichment analysis of genes differentially expressed (DEX) in MC cells (adjusted P-value < 0.05) compared to all other cell types in the middle turbinate. The analysis revealed significant enrichment in multiple biological processes related to development (**Supplemental Table 1**). Other noteworthy biological processes are those related to cell adhesion, cell migration, and mesenchymal cell maintenance (mesenchymal development, mesenchymal cell differentiation, regulation of epithelial to mesenchymal transition, positive regulation of epithelial to mesenchymal transition, epithelial to mesenchymal transition). While there is an enrichment of genes involved in neurogenesis (GO:0022008) and related processes (generation of neurons (GO:0048699), regulation of neuron projection development (GO:0010975), neuron projection development (GO:0031175), neuron projection morphogenesis (GO:0048812), etc.), processes involved in the development of some non-neuronal tissues and organs are also present.

Overall, the development of the human body is driven by the stem and progenitor cells, which divide, migrate, differentiate and shape different organs. There are several known sets of genes supporting stem cell properties, such as those described in embryonic, neural, intestinal, adipose-derived stem cells, mesenchymal stem cells in different tissues, and cancer stem cells.

Finally, we compared single cell gene expression data between CNON cells (CNON-CTRL and CNON-SCZ) to assess potential usability of scRNA-seq to study schizophrenia. CNON-SCZ sample was selected for this study because cells from CNON-SCZ have a low growth rate and extreme cell cycle eigengene value in our previous study (20). It was located on the periphery of PCA1/PCA2 map based on all or only DEX genes, suggesting that this sample may reveal schizophrenia-specific differences in expression profiles in single cell data despite the high genetic and transcriptomic heterogeneity of both CNON samples. Most of DEX genes identified in our previous study had low expression, with only 21 of them with transcripts per million transcripts (TPM) more than 10, and only 5 genes with TPM>100. In single cells, we found 15 DEX genes with detected expression in more than 50% of cells. Eleven of them showed difference in expression between control and SCZ sample of more than 50%, all of them in the same direction as it was assumed by DEX analysis in bulk RNA-seq study. We initially assumed that these alterations in gene expression could explain the dramatic reduction in proliferation rate, but single cell data revealed a different picture. The percentage of cells in G2-M and S cycle stage in the slow-growing CNON-SCZ sample was higher than in the fast-growing CNON-CTRL. Instead, we found that a substantially larger fraction of cells from SCZ sample was associated with a cluster of cells with elevated level of mitochondrial gene expression, often characteristic of apoptotic processes (**Table 3**). Given that apoptosis is relatively rapid process (21), slow growth of SCZ sample may potentially be explained by a higher apoptosis rate.

**Table 3.** Number and percentage of cells in G0, G1, S, G2/M and apoptotic clusters in CNON-CTRL and CNON-SCZ cell cultures.

|  | CNON-CTRL |  | CNON-SCZ |  |
| --- | --- | --- | --- | --- |
|  | Number of Cells | Percentage of Cells | Number of Cells | Percentage of Cells |
| G0 | 10854 | 88.7 | 2553 | 77.3 |
| G1 | 634 | 5.2 | 348 | 10.5 |
| S | 556 | 4.5 | 242 | 7.3 |
| G2-M | 143 | 1.2 | 106 | 3.2 |
| Apoptotic | 47 | 0.4 | 54 | 1.6 |

## Discussion

Cells in CNON lack prominent markers of pluripotency, such as *POU5F1* and *NANOG*, which are the hallmarks of embryonic stem cells and iPSCs. *PROM1* (CD133), a marker used for the purification of neural stem cells (22), is also not expressed in CNON. Currently, the literature on neural progenitor cells in the brain is less well described. One of the better-known types is radial glia (a cell that differentiates into outer radial glia and ventricular radial glia at later stages of brain development), which can directly differentiate into neurons or produce intermediate progenitor cells (IPCs). However, the expression profile of CNON does not fit into the radial glial gene expression pattern; in particular, CNON does not express radial glial markers *SOX2, HES5, PAX6*, or *GFAP*. CNON also does not express *EOMES*, a marker of IPCs, which play an important role in producing neurons after gestation week 8 (~PFA 6 weeks).

Neuroepithelium such as ON and epithelium of middle turbinate originates mostly from ectoderm, while cells from the MC cluster express multiple markers of mesenchymal cells, or mesenchymal stem cells, which are multipotent cells of mesodermal origin (**Table 4**). Precursors of these mesenchymal cells likely originate from neural crest, from which they migrate to different locations of the fetus to establish mesenchymal cell populations in bone marrow (23), adipose tissue (24), oral mucosa lamina propria (25), and several other locations (see (26) for review), including the brain (27). Expectedly, cluster 9 of embryonic brain sample CS14_3 also expressed multiple mesenchymal markers.

**Table 4.**
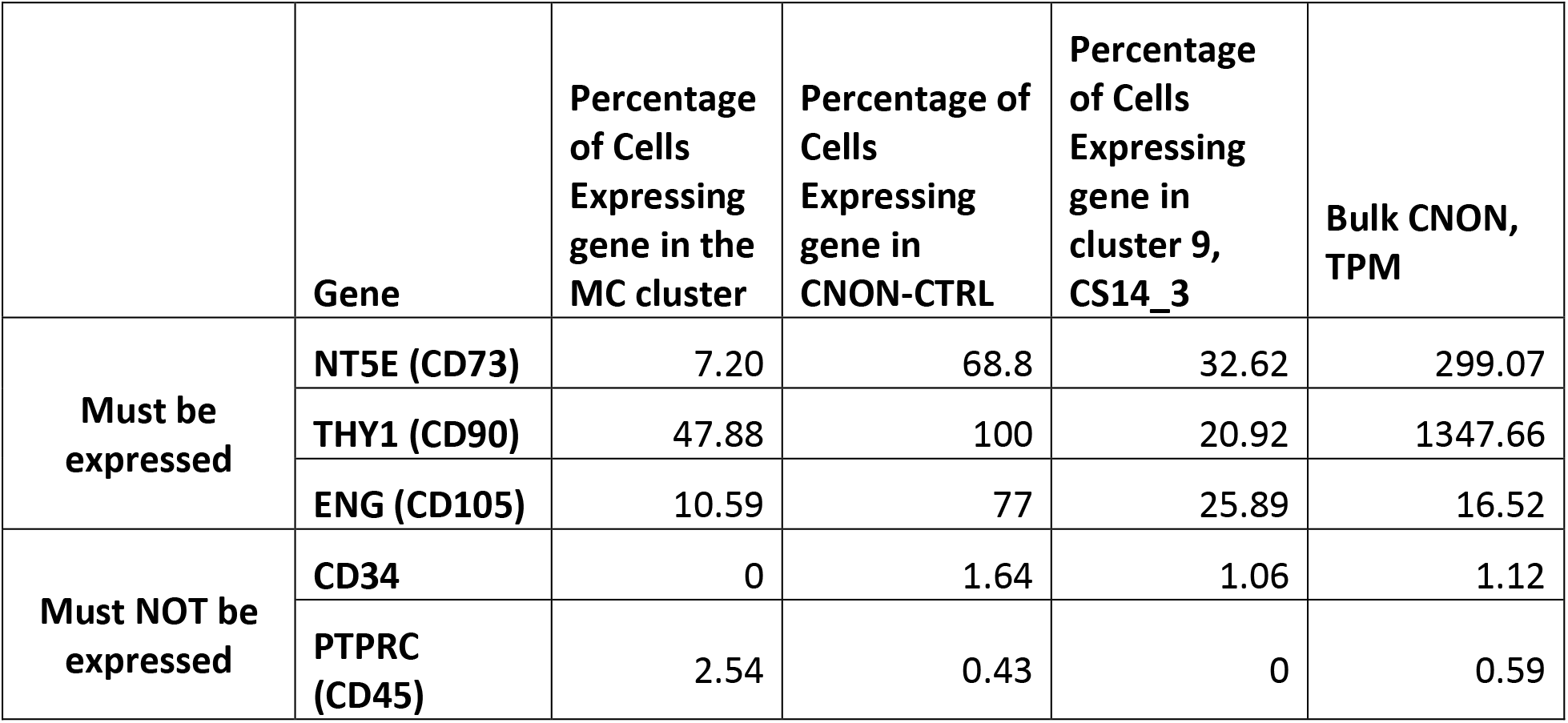
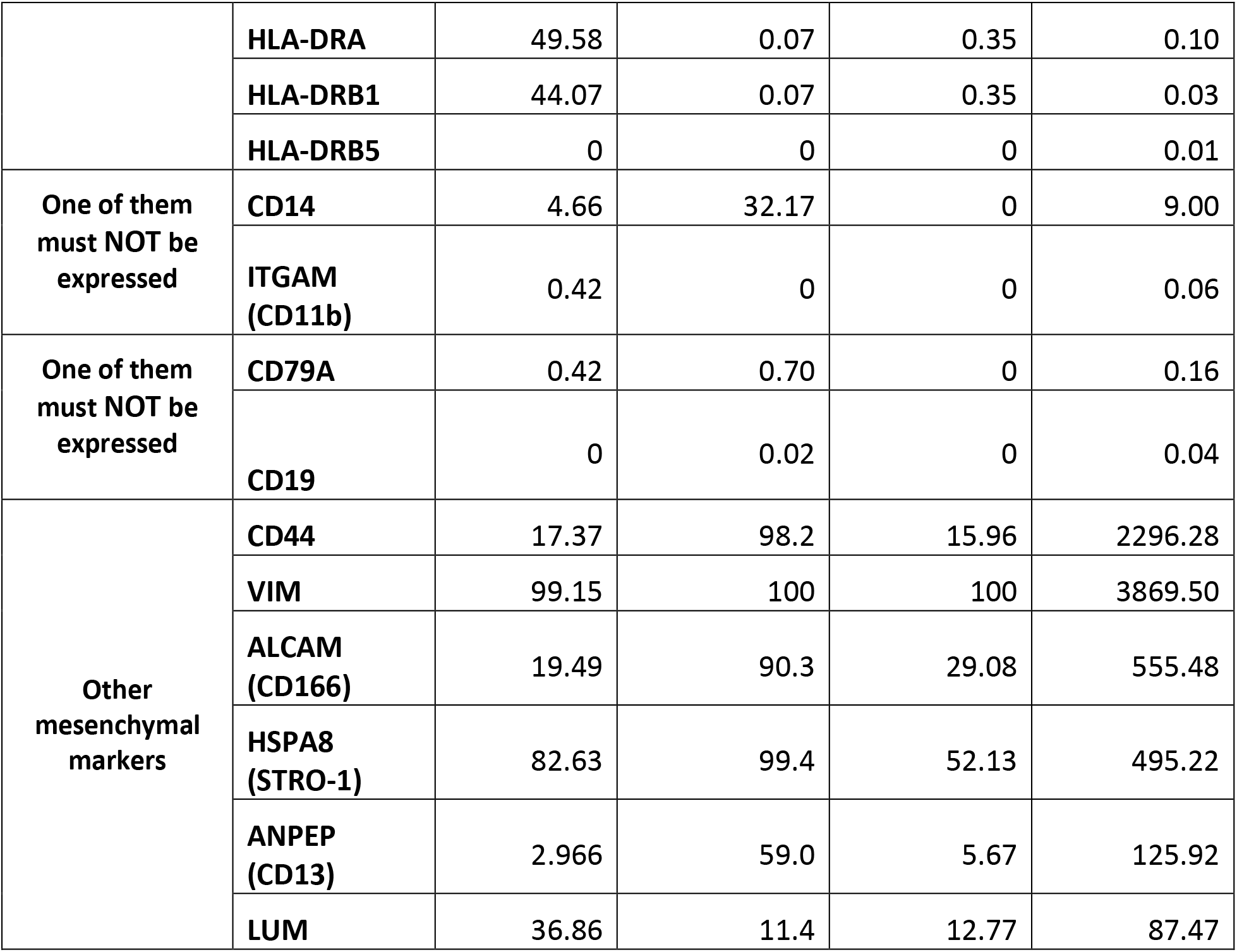
Expression of mesenchymal markers in the MC cluster (MT-CTRL), CNON (CNON-CTRL) and cluster 9 of embryonic brain (CS14_3, (19)) and bulk CNON (average Transcripts per Million transcripts from 255 CNON samples). According to the Mesenchymal and Tissue Stem Cell Committee of the International Society for Cellular Therapy (31), “MSC must express CD105, CD73 and CD90, and lack expression of CD45, CD34, CD14 or CD11b, CD79a or CD19 and HLA-DR surface molecules”. Other prominent markers of mesenchymal cells are also included in the table.

Despite the characteristic similarities in the expression of specific marker genes, mesenchymal cells from different tissues may differ in the expression of some genes and their ability to differentiate into particular cell types in a specific environment. For example, mesenchymal-like stem cells residing in the olfactory mucosa demonstrated the promyelination effect on oligodendrocyte precursor cell, while similar mesenchymal cells derived from bone marrow do not enhance myelination (28). Distinctions between MSC from different tissues define their specific use in regenerative medicine (29,30).

Mesenchymal and Tissue Stem Cell Committee of the International Society for Cellular Therapy has developed a set of minimum criteria for defining multipotent mesenchymal stromal cells: **(a)** on the expression of certain proteins/genes, **(b)** the ability to adhere to plastic and **(c)** differentiate *in vitro* into osteoblasts, adipocytes, chondroblasts (31). CNON satisfied the first two criteria of expressing specific genes and their ability to adhere to plastic (**Figure 2b**), but we have yet to investigate the multipotency of CNON.

**Figure 2.**
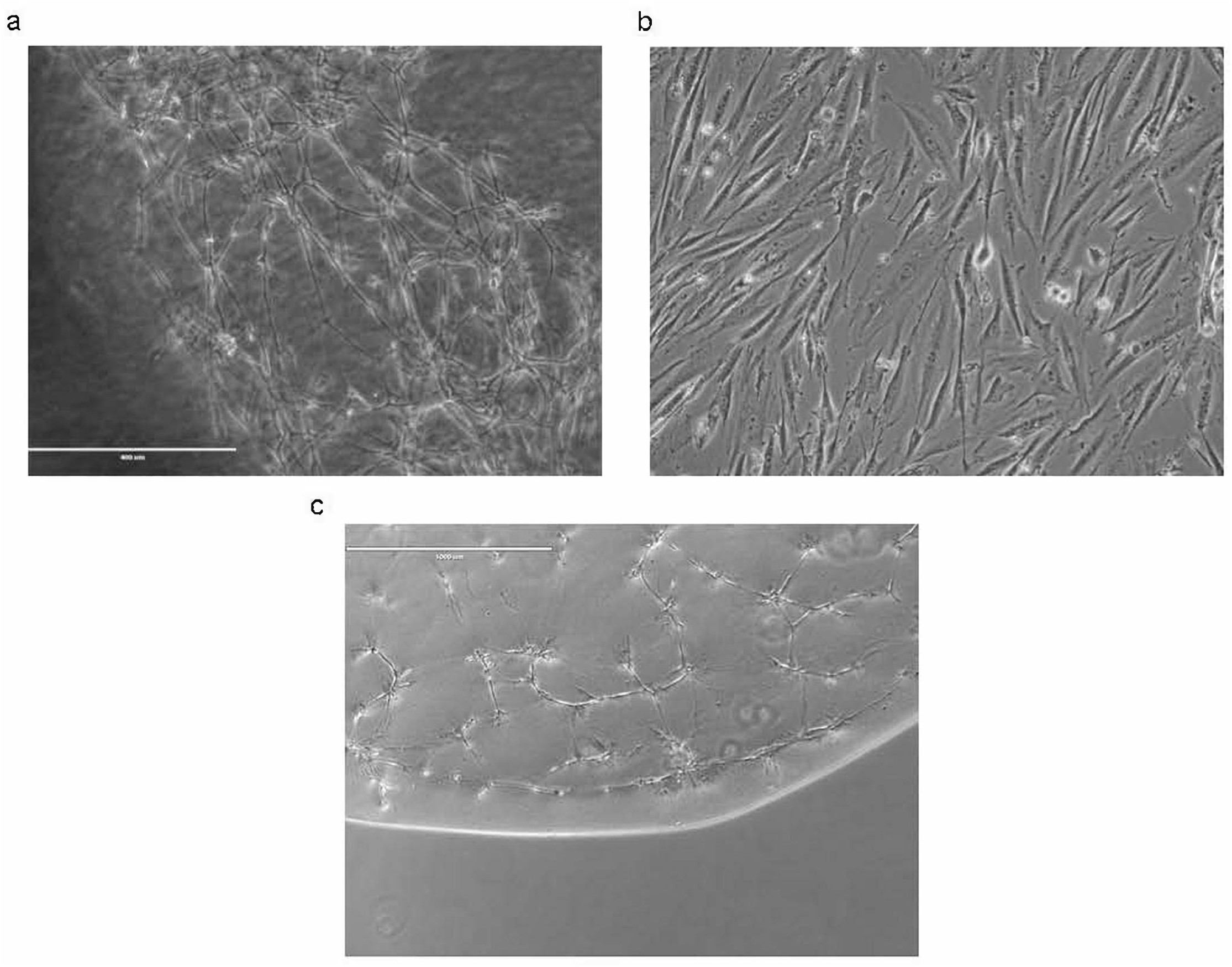
Microscopic images of CNON cell (A) outgrowing from biopsy piece in Matrigel, (B) growing in 2D culture, and (C) growing in Matrigel after culturing in 2D for several passages.

In previous studies, WNT signaling has been shown to play an important role in controlling both the maintenance and differentiation of mesenchymal stem cells (32–35). The CNON expression profile shows a robust expression of Wnt5A, Wnt5B and Notch2 ligands, accompanied by genes involved in corresponding signaling pathways. For example, genes for frizzled receptors *FZD2* and *FZD7*, co-receptors *ROR2, LPR5,LPR6*, and secreted frizzled receptors *SFRP1* and *SFRP2*, as well as *CTNNB1* (β-catenin), are robustly expressed in CNON to support both canonical and non-canonical signaling (**Table 5**). Similarly, *JAG1*, presenilins *PSEN1* and *PSEN2, ADAMS17, PSENEN*, and *APH1A* (γ-secretase subunits) are also expressed in CNON (**Table 5**). This suggests that CNON can regulate differentiation and function in an autocrine or paracrine manner.

**Table 5.**
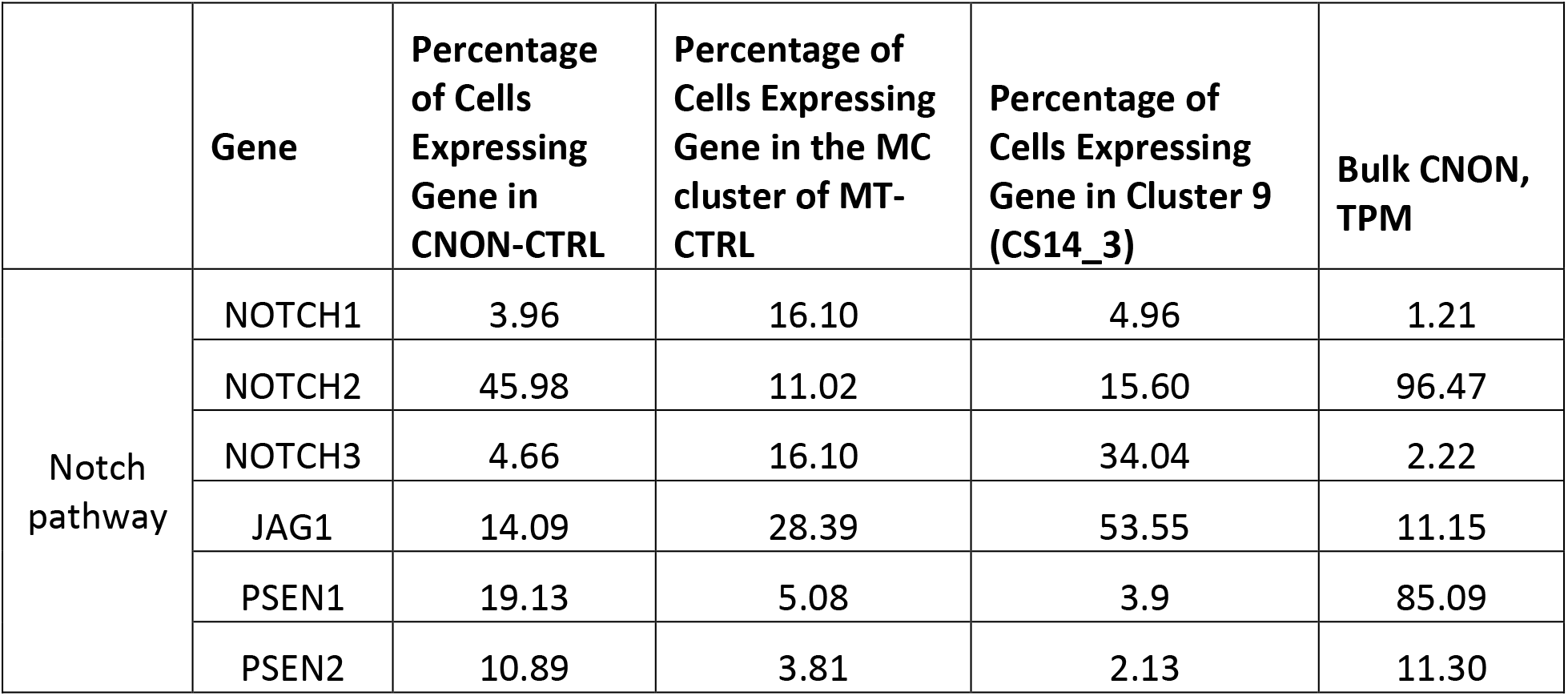
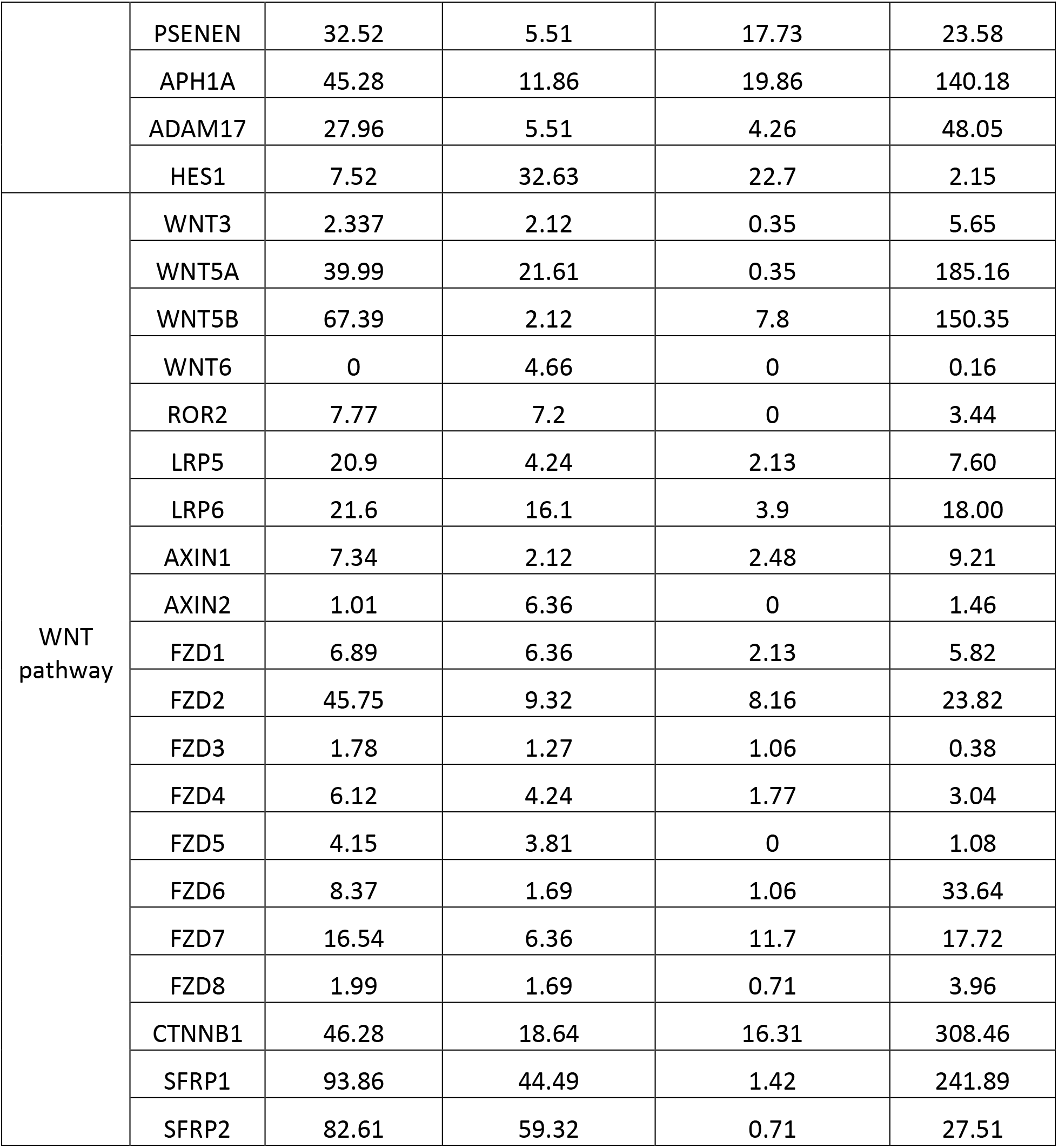
Expression of Wnt- and Notch-pathways genes in CNON (SEP036), the MC cluster of MT (SEP310), cluster 9 (CS14_3) of embryonic brain and bulk CNON (average Transcripts per Million transcripts from 255 CNON samples).

|  | Gene | Percentage of Cells Expressing Gene in CNON-CTRL | Percentage of Cells Expressing Gene in the MC cluster of MT-CTRL | Percentage of Cells Expressing Gene in Cluster 9 (CS14_3) | Bulk CNON, TPM |
| --- | --- | --- | --- | --- | --- |
| Notch pathway | NOTCH1 | 3.96 | 16.10 | 4.96 | 1.21 |
|  | NOTCH2 | 45.98 | 11.02 | 15.60 | 96.47 |
|  | NOTCH3 | 4.66 | 16.10 | 34.04 | 2.22 |
|  | JAG1 | 14.09 | 28.39 | 53.55 | 11.15 |
|  | PSEN1 | 19.13 | 5.08 | 3.9 | 85.09 |
|  | PSEN2 | 10.89 | 3.81 | 2.13 | 11.30 |
|  | PSENEN | 32.52 | 5.51 | 17.73 | 23.58 |
|  | APH1A | 45.28 | 11.86 | 19.86 | 140.18 |
|  | ADAM17 | 27.96 | 5.51 | 4.26 | 48.05 |
|  | HES1 | 7.52 | 32.63 | 22.7 | 2.15 |
| WNT<br>pathway | WNT3 | 2.337 | 2.12 | 0.35 | 5.65 |
|  | WNT5A | 39.99 | 21.61 | 0.35 | 185.16 |
|  | WNT5B | 67.39 | 2.12 | 7.8 | 150.35 |
|  | WNT6 | 0 | 4.66 | 0 | 0.16 |
|  | ROR2 | 7.77 | 7.2 | 0 | 3.44 |
|  | LRP5 | 20.9 | 4.24 | 2.13 | 7.60 |
|  | LRP6 | 21.6 | 16.1 | 3.9 | 18.00 |
|  | AXIN1 | 7.34 | 2.12 | 2.48 | 9.21 |
|  | AXIN2 | 1.01 | 6.36 | 0 | 1.46 |
|  | FZD1 | 6.89 | 6.36 | 2.13 | 5.82 |
|  | FZD2 | 45.75 | 9.32 | 8.16 | 23.82 |
|  | FZD3 | 1.78 | 1.27 | 1.06 | 0.38 |
|  | FZD4 | 6.12 | 4.24 | 1.77 | 3.04 |
|  | FZD5 | 4.15 | 3.81 | 0 | 1.08 |
|  | FZD6 | 8.37 | 1.69 | 1.06 | 33.64 |
|  | FZD7 | 16.54 | 6.36 | 11.7 | 17.72 |
|  | FZD8 | 1.99 | 1.69 | 0.71 | 3.96 |
|  | CTNNB1 | 46.28 | 18.64 | 16.31 | 308.46 |
|  | SFRP1 | 93.86 | 44.49 | 1.42 | 241.89 |
|  | SFRP2 | 82.61 | 59.32 | 0.71 | 27.51 |

Other studies also reported that cells with mesenchymal properties in the ON (ecto-mesenchymal stem cells) (29) and were able to culture them *in vitro* albeit using different cell culture method (36). As we demonstrated in *Results* section, single cell data from olfactory neuroepithelium (1) shows large group of cells with expression profile corresponding to CNON, which express multiple mesenchymal markers. Recent single cell study of cell cultures derived from olfactory mucosa also showed a large group of cells referred as “fibroblast/stromal”. However, at the time of writing, we are unable to gain access to the data to assess expression of mesenchymal markers. Notably, the study did not utilize selection for cells penetrating Matrigel, and resulting culture consists of multiple cell types, which were described as fibroblast/stromal, GBC and myofibroblasts (37).

Another study suggests that mesenchymal cells from olfactory mucosa possess multipotency and can form neurospheres different from those produced by horizontal epithelial global cells (38).

Therefore, the similarity of mesenchymal properties between these cells and CNON suggests that CNON is likely a derivative of ecto-mesenchymal stem cells of ON. CNON is a single cell type due to the specific way of developing cell cultures, while other methods without using Matrigel for cell type selection resulted in at least three different cell types growing in culture (37).

It should be noted that CNON have a more pronounced gene expression profile of mesenchymal cell markers compared to MC cluster of MT or cluster 9 from embryonic brain sample. For example, cells from MC in MTs express HLA-DR genes but their expression is halted when culturing them *in vitro* under our conditions. It is not unexpected, as the definition of mesenchymal stem cells is based on assays of cells in culture, and for a long time there was a hypothesis that MSC is an artifact of culturing cells *in vitro* (as discussed, for example, in (39)). We attribute it to plasticity of mesenchymal cells, changing phenotype and gene expression according to environment.

Indeed, cells in MC cluster in turbinates are not dividing; in Matrigel they change phenotype, divide and migrate (**Figure 2A**) and then change their phenotype again to classical mesenchymal phenotype when grown in 2D (**Figure 2b**). However, when placed inside Matrigel after multiple passages in 2D, cells reverted to a phenotype with multiple elongated branches and organizing complex interconnected cell structures resembling a neuronal network (**Figure 2c**).

Similar complex cellular structures have been previously observed when human mesenchymal stromal cells derived from bone marrow grew in diluted Matrigel, which the authors attributed to formation of capillary network (40). It may indicate versatility and plasticity of mesenchymal cells exploiting similar architectural solutions for different biological purposes.

The role of cells with mesenchymal gene expression signatures at the early stages of development of the brain is not clear. Many noticed similarities of MSC with pericytes and suggest the involvement of MSC in making a brain barrier and vascular systems. There is also an appreciation of their paracrine function in a neurovascular niche context involving in orchestrating the complex development of brain structures (27). Ability of MSC to differentiate into neurons *in vitro* and engrafted in the brain also suggest involvement of these cells in neurogenesis. Transplantation of MSC in brain lesion models showed therapeutic effects (41), although it is not yet clear if it is caused by MSC differentiation into neurons or by a paracrine effect or both. Anyway, the important role of MSC in the brain is well recognized, and the hypothesis that alterations in gene expression in MSC cause SCZ is well justified.

The role of mesenchymal cells in turbinates is not known. Although they have a capability to differentiate into neurons, it is not clear if they realize this potential in olfactory mucosa either during development or injuries or regular replenishment of olfactory neurons. The striking similarity of CNON to cells in embryonic brain is probably a reflection of similarities of MSC among tissues. MT belongs to the respiratory system, where the role of mesenchymal cells is quite pronounced. Mesenchymal cells are involved in lung development and responsible for homeostasis and tissue repair in the lung (42). If alterations in gene expression of mesenchymal cells contribute to the etiology of schizophrenia, we should expect that the same changes in MSC properties can affect other organs with substantial presence of MSC. Indeed, a comorbidity between SCZ and lung diseases have been reported, and in both directions: schizophrenia was found to be associated with impaired lung function (43), and patients with COPD have 10 times higher risk of psychiatric comorbidities (44). There are reports that olfactory deficits\ known to be prevalent in SCZ are also found in most COPD patients (45). We hypothesize that alterations in the properties of mesenchymal cells may contribute to a range of “mesenchymal” disorders, which include some subtypes of schizophrenia and lung diseases.

Our findings made on analysis of CNON that WNT signaling and regulation of WNT production (and specifically WNT5A), are involved in etiology of SCZ (19) fits this hypothesis well. These pathways are central for self-renewal and differentiation of MSC (33), and they are also play a critical role in lung development (46), tissue regeneration (47) and etiology of COPD (48). Thus, alterations in these pathways could be one of common mechanisms of “mesenchymal” disorders.

## Conclusion

Using scRNA-seq we confirmed that CNON cells originate from a single cell type of middle turbinate or olfactory neuroepithelium. The expression profile of CNON closely matches those of mesenchymal stem cells. Although we have not tested multipotency of these cells in this study, other studies suggest that mesenchymal cells of olfactory mucosa are able to differentiate into multiple lineages, including neurons.

Our analysis of gene expression in embryonic brain (2) identified a cell type that closely matches to CNON cells by expression profiling. Cell type homogeneity of CNON, stability of their expression profiles in cell culture during multiple passages, and high similarity to one of cell type in the embryonic brain providing support that CNON is a promising cellular model of neurodevelopmental disorders.

## Author Contributions

VSKT performed most of data analysis and made substantial contribution to manuscript preparation. MB performed biopsies. MB, FM, and RF provided expertise in respiratory epithelium physiology and cell biology. GS worked with cell cultures, performed single cells library preparation and sequencing. JK contributed to discussion and helped with organisation of the study. CP and MP organized recruitment of participants and contributed to discussion. OE designed and supervised all stages of the project, performed data analysis with VSTG and contributed to manuscript preparation.

## 1 Data Availability Statement

The data discussed in this publication have been deposited in NCBI’s Gene Expression Omnibus (49) and are accessible through GEO Series accession number GSE219165 (https://www.ncbi.nlm.nih.gov/geo/query/acc.cgi?acc=GSE219165). Original R scripts are available from the corresponding author, VSKT, upon reasonable request.

