## Supplemental Table 1 for "Cultured Mesenchymal Cells from Nasal Turbinate as a Cellular Model of the Neurodevelopmental Component of Schizophrenia Etiology"

**Supplemental Table 1.** Gene ontology enrichment analysis of genes differentially expressed in Mesenchymal cells cluster (adjusted P-value < 0.05) in comparison to all other cell types in the middle turbinate.

| GO biological process complete | fold Enrichment | P-value | P.adjust |
| --- | --- | --- | --- |
| regulation of neuron projection development (GO:0010975) | 5.21 | 1.59E-04 | 1.84E-02 |
| neuron projection development (GO:0031175) | 3.86 | 5.52E-04 | 4.16E-02 |
| neurogenesis (GO:0022008) | 3.6 | 6.93E-06 | 2.13E-03 |
| neuron development (GO:0048666) | 3.44 | 6.43E-04 | 4.63E-02 |
| generation of neurons (GO:0048699) | 3.32 | 1.28E-04 | 1.57E-02 |
| neuron differentiation (GO:0030182) | 3.27 | 2.80E-04 | 2.59E-02 |
| cell-matrix adhesion (GO:0007160) | 10.74 | 1.21E-04 | 1.50E-02 |
| cell-substrate adhesion (GO:0031589) | 8.97 | 6.48E-05 | 1.02E-02 |
| negative regulation of cell adhesion (GO:0007162) | 7 | 7.16E-05 | 1.05E-02 |
| cell adhesion (GO:0007155) | 5.39 | 4.06E-09 | 7.96E-06 |
| regulation of cell adhesion (GO:0030155) | 4.04 | 8.41E-05 | 1.16E-02 |
| tissue migration (GO:0090130) | 10.36 | 6.86E-04 | 4.89E-02 |
| ameboidal-type cell migration (GO:0001667) | 8.66 | 7.84E-05 | 1.10E-02 |
| negative regulation of cell migration (GO:0030336) | 7.05 | 6.86E-05 | 1.02E-02 |
| positive regulation of cell migration (GO:0030335) | 5.45 | 1.56E-05 | 3.45E-03 |
| cell migration (GO:0016477) | 4.82 | 4.12E-07 | 2.81E-04 |
| regulation of cell migration (GO:0030334) | 4.37 | 3.28E-06 | 1.25E-03 |
